# Visual feature tuning properties of stimulus-driven saccadic inhibition in macaque monkeys

**DOI:** 10.1101/2023.07.25.550484

**Authors:** Fatemeh Khademi, Tong Zhang, Matthias P. Baumann, Antimo Buonocore, Tatiana Malevich, Yue Yu, Ziad M. Hafed

## Abstract

Saccadic inhibition refers to a short-latency transient cessation of saccade generation after visual sensory transients. This oculomotor phenomenon occurs with a latency that is consistent with a rapid influence of sensory responses, such as stimulus-induced visual bursts, on oculomotor control circuitry. However, the neural mechanisms underlying saccadic inhibition are not well understood. Here, we exploited the fact that macaque monkeys experience robust saccadic inhibition to test the hypothesis that inhibition time and strength exhibit systematic visual feature tuning properties to a multitude of visual feature dimensions commonly used in vision science. We measured saccades in three monkeys actively controlling their gaze on a target, and we presented visual onset events at random times. Across six experiments, the visual onsets tested size, spatial frequency, contrast, motion direction, and motion speed dependencies of saccadic inhibition. We also investigated how inhibition might depend on the behavioral relevance of the appearing stimuli. We found that saccadic inhibition starts earlier, and is stronger, for large stimuli of low spatial frequencies and high contrasts. Moreover, saccadic inhibition timing depends on motion direction, with earlier inhibition systematically occurring for horizontally than for vertically drifting gratings. On the other hand, saccadic inhibition is stronger for faster motions, and when the appearing stimuli are subsequently foveated. Besides documenting a range of feature tuning dimensions of saccadic inhibition on the properties of exogenous visual stimuli, our results establish macaque monkeys as an ideal model system for unraveling the neural mechanisms underlying a highly ubiquitous oculomotor phenomenon in visual neuroscience.

**New and noteworthy:** Visual onsets dramatically reduce saccade generation likelihood with very short latencies. Such latencies suggest that stimulus-induced visual responses, normally jumpstarting perceptual and scene analysis processes, can also directly impact the decision of whether to generate saccades or not, causing saccadic inhibition. Consistent with this, we found that changing the appearance of the visual onsets systematically alters the properties of saccadic inhibition. These results constrain neurally-inspired models of coordination between saccade generation and exogenous sensory stimulation.

## Introduction

Saccadic inhibition is an inevitable consequence of exogenous visual sensory stimulation (1). In this phenomenon, the appearance of a visual stimulus, no matter how brief, is associated with an almost-complete cessation of saccade generation, and this cessation occurs with express latencies of less than 90-100 ms from stimulus onset (2–8). This conjunction of an early motor effect and a sensory origin driving it would suggest that saccadic inhibition reflects the arrival of visual sensory signals at the final oculomotor control circuitry relatively rapidly. Consistent with this, some studies in humans have demonstrated that saccadic inhibition depends on the contrast of the appearing visual stimuli (9–11); this reinforces the notion that saccadic inhibition can reflect visual sensory feature tuning properties somewhere late in the visual-motor hierarchy (1). Moreover, stimulus size exhibits a modulatory effect on the latency and strength of saccadic inhibition (6, 7). Exploiting the fact that saccadic inhibition and related smooth eye velocity modulations also occur during smooth pursuit eye movements (12–14), yet other studies have shown a potential dependence on spatial frequency of the inhibitory oculomotor processes associated with saccadic inhibition (15).

Despite the fact that monkeys, constituting a highly suitable animal model for investigating neural mechanisms, also show robust saccadic inhibition, whether in controlled fixation tasks (16, 17) or in free viewing paradigms (18, 19), the neural mechanisms driving saccadic inhibition remain elusive (1, 20). Earlier models have suggested that lateral inhibition in sensory-motor structures like the superior colliculus and frontal eye fields might play a role in this phenomenon (21–24). However, neither inactivation of the superior colliculus (25) nor the frontal eye fields (26) alters saccadic inhibition in any meaningful way. Moreover, the detailed feature tuning properties of saccadic inhibition in monkeys have not yet been fully documented. We recently showed that saccadic inhibition latency (and associated movement vector modulations) in macaque monkeys depends on the luminance polarity of the visual onsets (dark versus bright contrasts) as well as on whether the onsets were of a small spot or of a large full-screen flash (27). This leaves a great deal more to desire: the use of monkeys to study the neural mechanisms underlying saccadic inhibition requires much further characterization of the visual feature tuning properties of this highly ubiquitous phenomenon in these animals.

In this article, we document a series of dependencies of saccadic inhibition in rhesus macaque monkeys on different visual feature dimensions. These feature dimensions include stimulus size, spatial frequency, contrast, motion direction, and motion speed. We also contrast saccadic inhibition when different forms of gaze orienting behaviors are triggered by the visual onsets. We find that saccadic inhibition exhibits a primarily low-pass frequency tuning characteristic, occurring earlier for low than high spatial frequency stimulus onsets. We also find that the inhibition starts earlier for high contrast stimuli, as well as for horizontal versus vertical motion directions. Saccadic inhibition also starts earlier for large rather than small visual stimuli. These results suggest that the nature of the visual sensory signals present in the final oculomotor control circuits mediating saccadic inhibition can be quite distinct from the visual feature tuning properties of brain areas, such as early cortical visual areas, that might instead serve other aspects of scene analysis; the oculomotor system possesses its own filtered representation of the visual environment (28).

## Materials and methods

### Experimental animals and ethical approvals

We collected data from three adult, male rhesus macaque monkeys (*macaca mulatta*) aged 7-14 years, and weighing 9.5-12.5 kg. All experiments were approved by ethics committees at the regional governmental offices of the city of Tübingen.

### Laboratory setup and animal procedures

The bulk of the data were collected in the same laboratory as that described in our earlier studies (29–31). Specifically, we used a CRT display spanning approximately 31 deg horizontally and 23 deg vertically. The display was approximately 72 cm in front of the animals, and it had a refresh rate of 85 or 120 Hz. The display was linearized and calibrated, and we used grayscale stimuli throughout the experiments. In some experiments in monkeys A and F, we used an LCD display with a refresh rate of 144 Hz (AOC AG273QX2700), which was also linearized and calibrated. Some of the behavioral tasks (e.g. dependence on the contrast of small, localized stimuli; see Results) were obtained by re-analyzing behavioral data from an earlier study (29).

Data acquisition and stimulus control were realized through our custom-made system based on PLDAPS (32). The system connected a DataPixx display control device (VPixx Technologies) with an OmniPlex neural data processor (Plexon), and the Psychophysics Toolbox (33–35).

The monkeys were prepared for the experiments earlier, since they also contributed to several earlier publications by our laboratory; for example, see refs. (29, 36). In the present purely behavioral experiments, we only measured eye movements using high performance eye tracking. To do so, we exploited an implantation of a scleral search coil that we had previously done in one eye of each monkey, and we used the magnetic induction technique to track eye position (37, 38). Naturally, additional follow-up neurophysiological experiments in these animals will use the knowledge generated here to try to better understand the neural mechanisms underlying saccadic inhibition. Head position was comfortably stabilized during the experiments by attaching a small head-holder device implanted on the skull with a reference point on the monkey chair.

### Experimental procedures

In each experiment, the monkeys fixated a central fixation spot (square of approximately 5.4 by 5.4 min arc) presented over a gray background. The fixation spot was either black or white (depending on the experiment and date it was run), and it was only white in the experiment in which the subjects were instructed to generate a foveating saccade towards the appearing peripheral stimulus (see Experiment 5 below). After an initial period of fixation, typically lasting between 500 and 1000 ms, a visual onset took place, which triggered saccadic inhibition. That is, we exploited the fact that microsaccades during fixation in monkeys continuously optimize eye position on the fixation spot (18, 39, 40); thus, they represent active oculomotor exploratory behavior on a miniature scale, which is fundamentally not different from free viewing (41). This is similar to human oculomotor behavior as well (42). Stimulus onsets of any kind robustly trigger an inhibition of these small saccades during active gaze control near the fixation spot, and this happens with a similar time course of saccadic inhibition to the case of free viewing saccades (16, 18). Thus, we characterized saccadic inhibition in this oculomotor context.

Across different experiments, we varied the type of visual onset that took place, as we explain in more detail next.

#### Experiment 1: Size tuning

During maintained fixation, a brief flash (∼12 or ∼7-8 ms duration) of a black circle centered on the fixation spot appeared. The circle had variable radius across trials from among eight possible values: 0.09, 0.18, 0.36, 0.72, 1.14, 2.28, 4.56, and 9.12 deg. Thus, we spanned a range of sizes from approximately the size of the fixation spot being gazed towards by the saccades (0.09 deg) to approximately the size of the full display (9.12 deg).

We typically ran this experiment in daily blocks of approximately 200-500 trials per session, and we collected a total of 7178, 9078, and 3103 trials in monkey A, F, and M, respectively. This resulted in a total of 628-1402 analyzed trials per condition per animal (after some exclusions, like when there were blinks around stimulus onset; see *Data analysis* below).

#### Experiment 2: Spatial frequency tuning

In this set of experiments, we presented a vertical sine wave grating of high contrast (100%). The grating remained on until trial end a few hundred milliseconds later (300 ms). The monkeys were required to maintain fixation on the visible fixation spot. Across trials, the grating could have one of five different spatial frequencies as follows: 0.5, 1, 2, 4, and 8 cycles/deg (cpd). The grating size was constrained by a square of 6 by 6 deg centered on the fixation spot. However, for some sessions in monkey F, the grating filled the entire display.

The results were the same for the different grating sizes (since 6 by 6 deg was already relatively large), so we combined them in our analyses. The phase of the grating was randomized across trials.

We typically ran this experiment in daily blocks of approximately 150-400 trials per session, and we collected a total of 2426 and 2032 trials in monkeys A and F, respectively. This resulted in a total of 380-487 analyzed trials per condition per animal.

#### Experiment 3: Contrast sensitivity with full-screen stimuli

In this set of experiments, the stimulus onset during active gaze fixation was a single display frame (∼12 or ∼7-8 ms) that was darker than the background (i.e. negative luminance polarity). This single-frame flash, which filled the entire display with a uniform luminance, could have the following contrast levels relative to the background (Weber contrast): 5%, 10%, 20%, 40%, and 80%.

We typically ran this experiment in daily blocks of approximately 200-600 trials per session. In total, we collected 4623, 4035, and 3946 trials in monkey A, F, and M, respectively. This resulted in a total of 760-1321 analyzed trials per condition per animal.

#### Experiment 4: Contrast sensitivity with small, localized stimuli

Here, we analyzed data from the fixation experiments of (29). That is, there was a stimulus onset during fixation consisting of a circle of 0.51 deg radius appearing somewhere on the display and staying on until trial end. The stimulus could have one of five different negative polarity (i.e. dark) Weber contrasts as follows: 5%, 10%, 20%, 50%, and 100%. We did not analyze the positive polarity (i.e. bright) contrasts from the previous study (29), because we wanted to compare saccadic inhibition to the task above with full-screen stimuli (but the results were generally similar).

We had a total of 3854 and 8551 analyzed trials from monkeys A and M, respectively, in this task. This resulted in 623-1692 trials per condition per animal.

#### Experiment 5: Contrast sensitivity with small, localized stimuli and visually-guided foveating movements towards them

In this case, we used a similar task to the one immediately above (Experiment 4), except that we removed the fixation spot as soon as the peripheral stimulus appeared (29). This allowed the monkeys to generate a foveating saccade towards the appearing stimulus immediately after the saccadic inhibition that was triggered by the stimulus onset was completed. Our goal here was to compare the inhibition properties when the appearing stimulus was oriented towards with a foveating eye movement, as opposed to being completely ignored. That is, we tested what happens when the appearing stimulus (which was outside of the range of ongoing eye movement target locations when it occurred) was either ignored (Experiment 4) or oriented towards (current experiment). The task was the same as the visually-guided saccade task described in (29).

We included a total of 1928, 3560, and 2474 trials from monkeys A, F, and M in our analyses of this task. This resulted in approximately 52-915 trials per condition per animal in our analyses. Note that in this experiment, not all contrasts were available as in Experiment 5.

Thus, the plots in Results only show data from the contrasts that were actually tested.

#### Experiment 6: Motion direction and speed

This experiment was similar to the spatial frequency tuning one above (Experiment 2) but with a constrained stimulus size (6 by 6 deg centered on the fixation spot location). In the current case, the grating presented was a drifting grating having one of eight equally spaced motion directions and one of two temporal frequencies (4 or 16 Hz; equivalent to 3.64 and 14.55 deg/s motion speeds, respectively). The spatial frequency was constant across all trials: 1.1 cycles/deg. At trial onset, the drifting grating appeared for 300 ms before the monkeys were rewarded for keeping their gaze near the central fixation spot.

We included a total of 3878, 1218, and 6327 trials from monkeys A, F, and M in our analyses of this task. This resulted in approximately 148-756 trials per motion direction (both speeds) per animal in the analyses.

### Data analysis

We detected all saccades using our established methods (43, 44). In all experiments, we included all saccades that happened in the peri-stimulus interval (regardless of their size), especially because we expected saccadic inhibition by stimulus onsets to affect all occurring movements in the monkeys (16). However, since the animals were engaged in fixation on a small target, the saccades were generally small anyway (e.g. median 18, 10, and 32 min arc in the pre-stimulus baseline fixation intervals of Experiment 6 in monkeys A, F, and M, respectively; similar values were observed in the other experiments).

In the orienting version of the contrast sensitivity task (Experiment 5), we also detected the foveating saccade towards the appearing stimulus. This allowed us to limit the upper temporal boundary for analyzing the timing of saccadic inhibition (see below for how we estimated saccadic inhibition timing). In other words, once a foveating saccade is generated, no subsequent saccadic inhibition could occur because the foveating saccade can only proceed after the oculomotor system has already been reset (1).

We excluded trials if there were blinks in the peri-stimulus interval that we were interested in analyzing (from -500 ms to +1000 ms relative to stimulus onset). We also excluded trials in which the monkeys broke their required gaze fixation state (either on the fixation spot or the foveated stimulus in the saccade task) before trial end. These were rare.

To compute saccade rate, we aggregated saccade onset times from all trials of a given condition and animal (we pooled data from the same condition across days of data collection in a given animal, but we always analyzed each monkey’s data separately). We then created arrays that were 0 at all times except for the time samples of saccade onsets (assigned a value of 1; 1000 Hz sampling rate). We then used a moving window of 50 ms, moving in steps of 1 ms, in which we counted the number of saccade onsets happening within the averaging window and within a given trial. This gave us a rate estimate per trial. We then averaged across all trials to obtain the average saccade rate curve of the particular condition. This approach is similar, in principle, to other standard saccade rate calculation approaches in the literature (11, 27). Subsequent analyses were made on the saccade rate curves that we obtained with this procedure.

Since saccadic inhibition happens very shortly after putative visual bursts in potential brain areas mediating the inhibition, we looked for hallmarks of feature tuning in the very initial phases of the stimulus-driven eye movement inhibition. To do this, we computed an estimate of the latency of the inhibition (called L_50_), and we related this latency to the different stimulus properties. Figure 1A describes the conceptual idea of the L_50_ measure, which we defined as done previously in the literature (3, 11, 45). Briefly, we first measured baseline saccade rate in the final 100 ms of fixation before stimulus onset in any given condition. We did this by averaging saccade rate over this 100 ms period and pooling across all trials of the condition (e.g. for all trials with 0.5 cpd in Experiment 2). We then estimated how much the rate dropped after stimulus onset during saccadic inhibition (i.e. the difference between the baseline rate and the minimum saccade rate after the stimulus onset). L_50_ was defined as the time point at which half of the rate drop during saccadic inhibition was achieved; the detailed robust estimate of this halfway drop is described exhaustively elsewhere (3, 11, 45). This measure is also conceptually similar to other estimates of saccadic inhibition timing (8). We then repeated this procedure for all other conditions.

**Figure 1.**
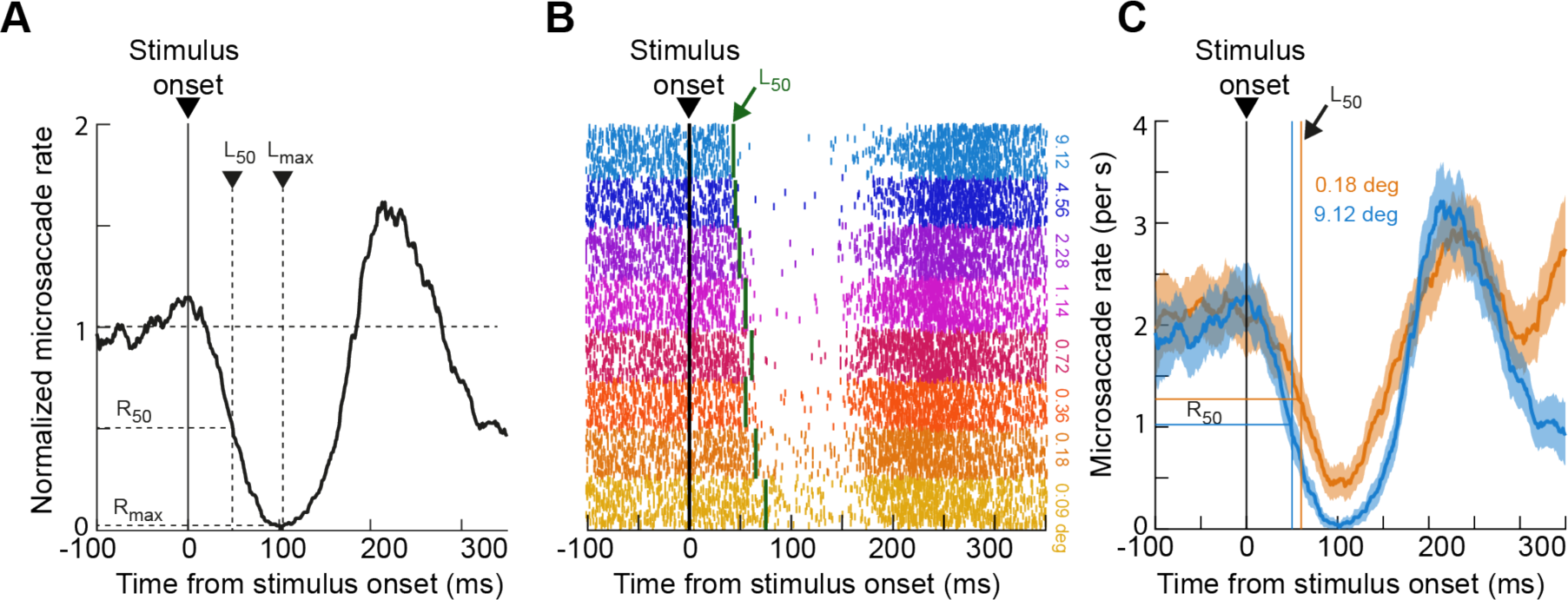
Relating saccadic inhibition to stimulus properties. **(A)** Example normalized saccade rate plot from one monkey and one condition. We were primarily interested in the time of saccadic inhibition, which we estimated via the L_50_ parameter described in the text; briefly, L_50_ indicates the time at which saccade rate dropped from baseline by half of the magnitude of its maximal drop caused by stimulus onset. We also reported R_50_, which is the raw saccade rate at the time of L_50_. **(B)** Example relationship between L_50_, saccadic inhibition, and stimulus properties from one animal and one experiment. The figure shows all saccades occurring around stimulus onset that monkey F generated during Experiment 1 (size tuning). Each row is a trial, and each tick mark is a saccade onset time. The trials were grouped according to the size of the appearing stimulus, and the vertical green lines indicate L_50_ estimates for each condition. As can be seen, L_50_ robustly indicated the timing of saccadic inhibition, which also clearly depended on stimulus appearance. **(C)** Example saccade rate from one monkey (A) and two conditions of Experiment 1. Error bars denote 95% confidence intervals. The x- and y-axis drop lines indicate the L_50_ and R_50_ values for each condition, respectively. As can be seen, saccadic inhibition timing reflected the change in stimulus property (in this case, size), also consistent with **B** in a second monkey. Figure 2 shows the full parametrization of size tuning of saccadic inhibition in all three monkeys.

Our L_50_ measure was a robust estimate of saccadic inhibition timing, as can be seen from Fig. 1B. This figure plots the raw saccade onset times of Experiment 1 from one example monkey (F). The saccades are graphed as raster plots with each row being a trial and each tick mark indicating saccade onset time relative to stimulus onset. Trials of the same type were grouped together and color-coded similarly for easier visualization (even though they were randomly interleaved during data collection). For each stimulus type, Fig. 1B also indicates the obtained estimate of L_50_. As can be seen, this measure was a robust estimate of saccadic inhibition timing.

Even though L_50_ was our parameter of primary interest in this study (given the above text and Fig. 1B), we also sometimes reported R_50_, which was simply the raw saccade rate (not- normalized to the baseline rate) at which L_50_ was reached (Fig. 1A). This allowed us to document general variability of microsaccade rate (whether in baseline or at the L_50_ time of saccadic inhibition) across individual monkeys. The calculation of R_50_ was again based on previously published methods (11, 45).

Note also that we were not interested in post-inhibition saccades (and how these saccades might depend on the visual stimulus properties). Post-inhibition saccades reflect reprogrammed movements after the inhibition (1, 40, 46), and they depend on frontal cortical activity (26, 47, 48); we were, instead, interested in the immediate impact on eye movements as revealed by L_50_. Nonetheless, for every experiment, we did plot example saccade rate curves that included the post-inhibition movements as well, for completeness (e.g. Fig. 1C).

Table 1 in the Appendix provides descriptive statistics of L_50_ and R_50_ in all experiments and all animals, as well as estimates of baseline saccade rates in each animal and the total number of trials analyzed per condition. To obtain estimates of 95% confidence intervals for each measure of L_50_ and R_50_ in Table 1, we used bootstrapping. Briefly, if a condition had N trials, we randomly selected N trials (with replacement) in a given bootstrap, and we calculated L_50_ and R_50_. We then repeated this process 1000 times. The 95% confidence intervals were the intervals encompassing the range between the 2.5th and 97.5th percentiles of all of the 1000 bootstrapped means. The obtained confidence intervals are also listed in Table 1 in the Appendix.

When documenting the potential influence of a visual feature (e.g. contrast) on saccadic inhibition time (L_50_), we also obtained the L_50_ measure for each condition and plotted it against the condition value (e.g. L_50_ versus contrast). For the size tuning, spatial frequency, and contrast manipulations, we often noticed that L_50_ (and sometimes R_50_) changed (either increased or decreased) with increasing stimulus size, spatial frequency, or contrast in an approximately logarithmic manner (see Results). Thus, we obtained a fit to a function of the form: L_50_ = a*log_10_(x) + b, where x is the parameter being varied in an experiment (e.g stimulus size or contrast) and a, b are the parameter fits. We included the fits in all relevant figures in Results, with indications of r^2^ values. We also used a similar approach for R_50_ plots, for completeness.

## Results

We characterized the timing of saccadic inhibition (L_50_; Materials and Methods) as a function of visual stimulus properties across a series of feature manipulations in three different animals (Fig. 1). We were motivated by the hypothesis that saccadic inhibition reflects the impact of short-latency stimulus-driven visual bursts on final oculomotor pathways (1). If so, then feature changes that are expected to alter visual responses (somewhere in the brain that is relevant for the inhibition) should also alter the time of saccadic inhibition. For example, in Fig. 1C, two different stimulus sizes from Experiment 1 resulted in two different timings of saccadic inhibition in an example monkey. Therefore, we adopted a descriptive approach in this study, documenting our observations on saccadic inhibition in multiple feature dimensions.

Our efforts across all experiments described below motivate a search (in macaque monkeys) for neural loci in the final oculomotor control circuitry, possibly in the brainstem, that would exhibit stimulus-driven visual bursts of neural activity matching the feature tuning properties of saccadic inhibition that we document below. This would mean that early sensory areas (such as retina, lateral geniculate nucleus, and primary visual cortex) relay rapid visual signals to visually-sensitive oculomotor areas, which might in turn reformat (28) these signals for specific use by the eye movement system, and for mediating the actual saccadic inhibition.

In the results below, besides saccadic inhibition timing (L_50_), we also documented our measures of R_50_ (Materials and Methods) because they roughly corresponded with the L_50_ modulations. Briefly, R_50_ describes the raw saccade rate at the L_50_ time. However, as stated above, we believe that the L_50_ modulations are the more meaningful ones, in general, since inhibition can be an all-or-none phenomenon in monkeys, especially for supra-threshold stimuli; this renders R_50_ closer to a floor effect for most stimulus features.

As also stated above, we additionally did not explicitly analyze post-inhibition saccades (besides plotting saccade rate curves to include their time ranges). This was so because such post-inhibition saccades reflect later processes (possibly also cognitively driven) (23, 49) that are needed to resume active oculomotor behavior after stimulus-driven interruption (also see the results of Experiment 5 below). Indeed, prior work has shown that these post- inhibition saccades may be governed by different underlying neural processes from those generating the more reflexive phenomenon of saccadic inhibition (26, 40, 46–48).

### Larger stimuli cause earlier saccadic inhibition

In our first experiment, we briefly presented a black circle centered on the fixation spot (Materials and Methods). Across trials, the circle could have one of eight different radii, ranging from 0.09 deg (approximately the size of the fixation spot) to 9.12 deg (approximately filling the whole display). We found that saccadic inhibition times roughly monotonically decreased with increasing stimulus size, as demonstrated in Fig. 2. This figure is organized as follows. For each animal (Fig. 2A for monkey A, Fig. 2D for monkey F, and Fig. 2G for monkey M), we first showed the saccade rate modulation time courses as computed in Fig. 1A, C. Here, each curve represents a different stimulus size that was presented. As can be seen, saccadic inhibition started earlier for larger onset sizes, and the dependence on size was roughly logarithmic. Specifically, Fig. 2B, E, H shows measures of L_50_ (our estimate of saccadic inhibition time; Fig. 1 and Materials and Methods) as a function of stimulus radius using a logarithmic x-axis. In all three animals, the data roughly followed a straight line (goodness of fits to a logarithmic curve are indicated in the respective figure panels). Thus, with a flash as little as 1-2 deg in radius, saccadic inhibition was already rendered even more robust than it was for smaller visual transients, and the effect eventually approached a plateau with even larger stimuli.

**Figure 2.**
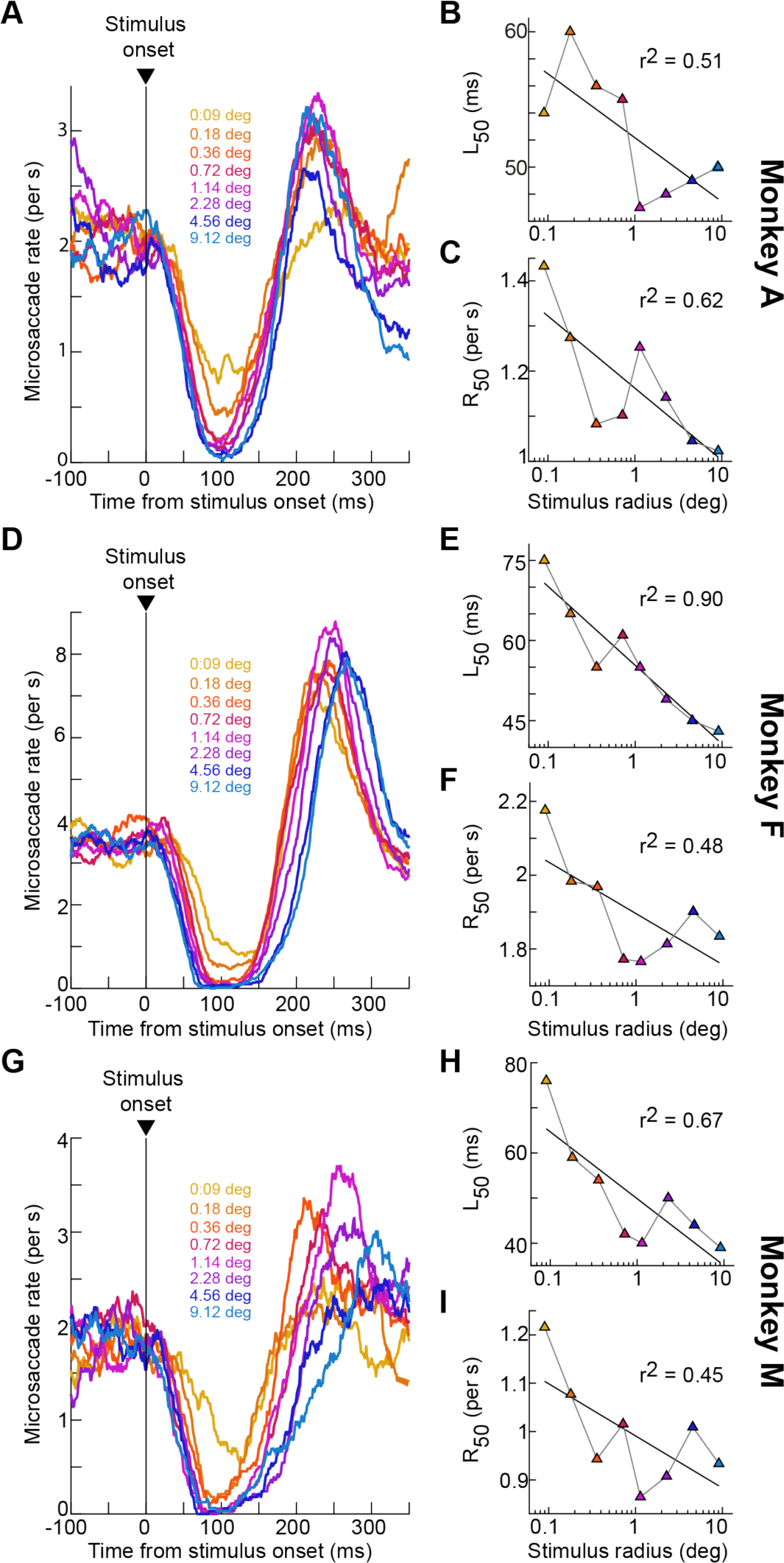
Earlier saccadic inhibition with larger stimuli. **(A)** Saccade rate curves relative to stimulus onset (like in Fig. 1) from monkey A in our size tuning experiment. Each colored curve corresponds to a stimulus radius as per the color-coded legend. Larger stimuli were associated with earlier and stronger saccadic inhibition. **(B)** A measure of saccadic inhibition time (L_50_) as a function of stimulus radius (Materials and Methods). Saccadic inhibition started earlier with larger stimuli, and the effect followed a roughly logarithmic relationship: the black line describes the fit to a logarithmic function (Materials and Methods) with the shown r^2^ value. **(C)** Similar to **B** but for a measure of saccadic inhibition strength (R_50_; Materials and Methods). Again, there was a stronger inhibition with larger stimulus sizes. **(D-F)** Similar observations from monkey F. **(G-I)** Similar observations from monkey M.

The results for R_50_ (the actual raw saccade rates at the time of L_50_; Materials and Methods) mimicked the above observations of L_50_, as can be seen from Fig. 2C, F, I. This is consistent with human observations (6, 7). Note, however, that L_50_ may be the more sensitive measure of stimulus-dependent changes in saccadic inhibition since saccade rate drops to almost zero (i.e. hits a floor effect) for most stimulus sizes (e.g. Fig. 2A, D, G). This is why our primary focus in this article, in general, was to document the L_50_ effects.

Thus, in rhesus macaque monkeys, saccadic inhibition shows a clear dependence on visual transient size, providing a clear homolog of human results with saccades in a different context (6, 7). This motivates using macaque monkeys to study the neurophysiological mechanisms underlying saccadic inhibition.

### High spatial frequencies are associated with delayed saccadic inhibition

We next turned our attention, in a second experiment, to the influences of spatial frequency on saccadic inhibition in rhesus macaque monkeys. Here, the monkeys fixated a central fixation spot while we presented a vertical sine wave grating of variable spatial frequency across trials. The grating stayed on the display until the monkeys were rewarded 300 ms later, and in some cases, it filled the whole display (Materials and Methods). Figure 3A shows the saccade rate curves of monkey A in this experiment. As can be seen, saccadic inhibition was systematically delayed with increasing spatial frequency of the appearing stimuli. This dependence was again roughly logarithmic, as can be seen from Fig. 3B and the associated logarithmic function fit (Materials and Methods). L_50_ in this animal was around 55 ms for 0.5 cpd gratings, but it was almost 85 ms for 8 cpd gratings. In this monkey, R_50_ did not systematically change as a function of spatial frequency (Fig. 3C). This is likely because the monkey’s pre-stimulus saccade rate was time varying (continuously decreasing) as a result of the animal systematically reducing its baseline saccade rate in anticipation of trial end; this time varying baseline added variability to our R_50_ measures.

**Figure 3.**
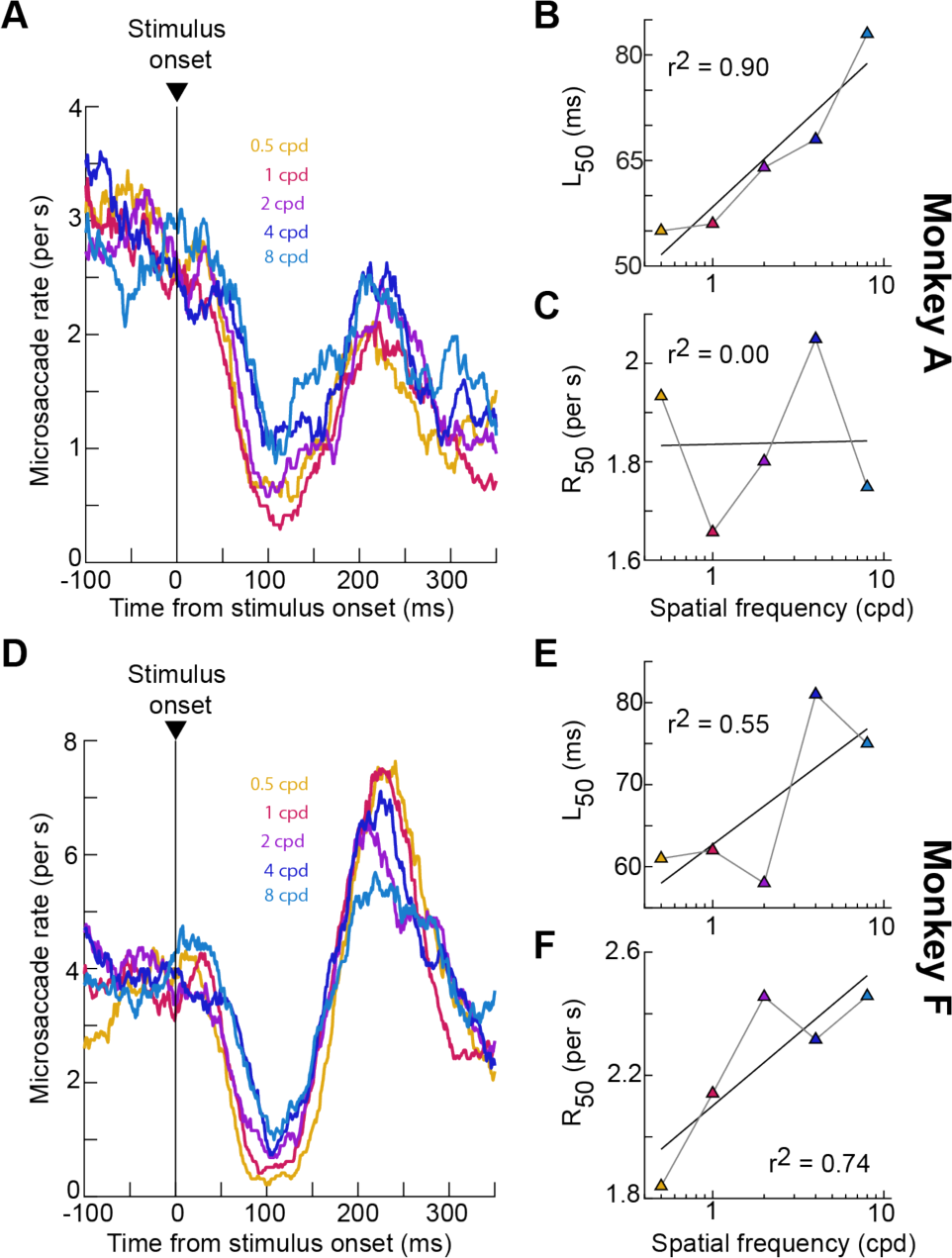
Earlier saccadic inhibition with lower spatial frequency stimuli. **(A)** Saccade rate curves from monkey A in the spatial frequency tuning experiment. Each curve now reflects saccade rate modulations for a stimulus onset of a given spatial frequency (indicated by the color-coded legend). There was earlier and stronger saccadic inhibition for the low spatial frequency stimulus onsets. **(B)** Inhibition time (L_50_) as a function of spatial frequency. This figure is formatted similarly to Fig. 2B, E, H. Inhibition time increased with a roughly logarithmic dependence as a function of increasing spatial frequency; the black line describes the best fitting logarithmic function equation (same as in Fig. 2) to the data (Materials and Methods). **(C)** Inhibition magnitude as assessed with R_50_ for the same data. Here, there was no clear relationship between R_50_ and spatial frequency (see text). **(D-F)** Similar analyses for monkey F. In this case, not only L_50_, but R_50_ also increased with stimulus spatial frequency.

In the second monkey that we tested with this task (monkey F), very similar observations were made for L_50_: the time of saccadic inhibition systematically increased with increasing spatial frequency (Fig. 3D, E). Since this monkey’s baseline (pre-stimulus) saccade rate was more constant than in monkey A, and also since the monkey’s minimum saccade rate during inhibition was markedly different from the baseline rate (Fig. 3D), the R_50_ measure also showed an increasing dependence on spatial frequency like for L_50_. Since R_50_ reflects the dynamic range of saccadic inhibition strength (Materials and Methods), this means that in addition to being later, saccadic inhibition was also weaker with higher spatial frequencies (minimum saccade rate during inhibition was higher).

Thus, saccadic inhibition in rhesus macaque monkeys shows a low-pass spatial frequency tuning characteristic. It is interesting that visual processing in the oculomotor system does also exhibit low-pass spatial frequency tuning properties (50, 51). This might suggest that as signals proceed from the retina and through the early visual system, the relevant visual response characteristics that might ultimately shape the feature tuning properties of saccadic inhibition can be different from the characteristics of early visual areas like primary visual cortex (which exhibits more band-pass spatial frequency tuning).

### Earlier saccadic inhibition with higher contrasts

Since previous human experiments demonstrated a dependence of saccadic inhibition on stimulus contrast (9–11), our next set of manipulations focused on this visual feature. The first such manipulation involved the onset of a full-screen flash of variable contrast across trials. The flash was of negative luminance polarity (darker than the background), and it occurred at a random time during fixation (Materials and Methods). In all three monkeys tested with this task, saccadic inhibition clearly occurred earlier for higher contrasts than for lower ones (Fig. 4A, D, G). Moreover, the time of L_50_ was again approximately logarithmically related to contrast level (Fig. 4B, E, H). Thus, consistent with humans, rhesus macaque monkeys show a dependence of saccadic inhibition timing on stimulus contrast. Our measures of R_50_ also behaved similarly to L_50_ (Fig. 4C, F, I), suggesting a larger drop in saccade likelihood at the time of peak saccadic inhibition for high contrast stimuli.

**Figure 4.**
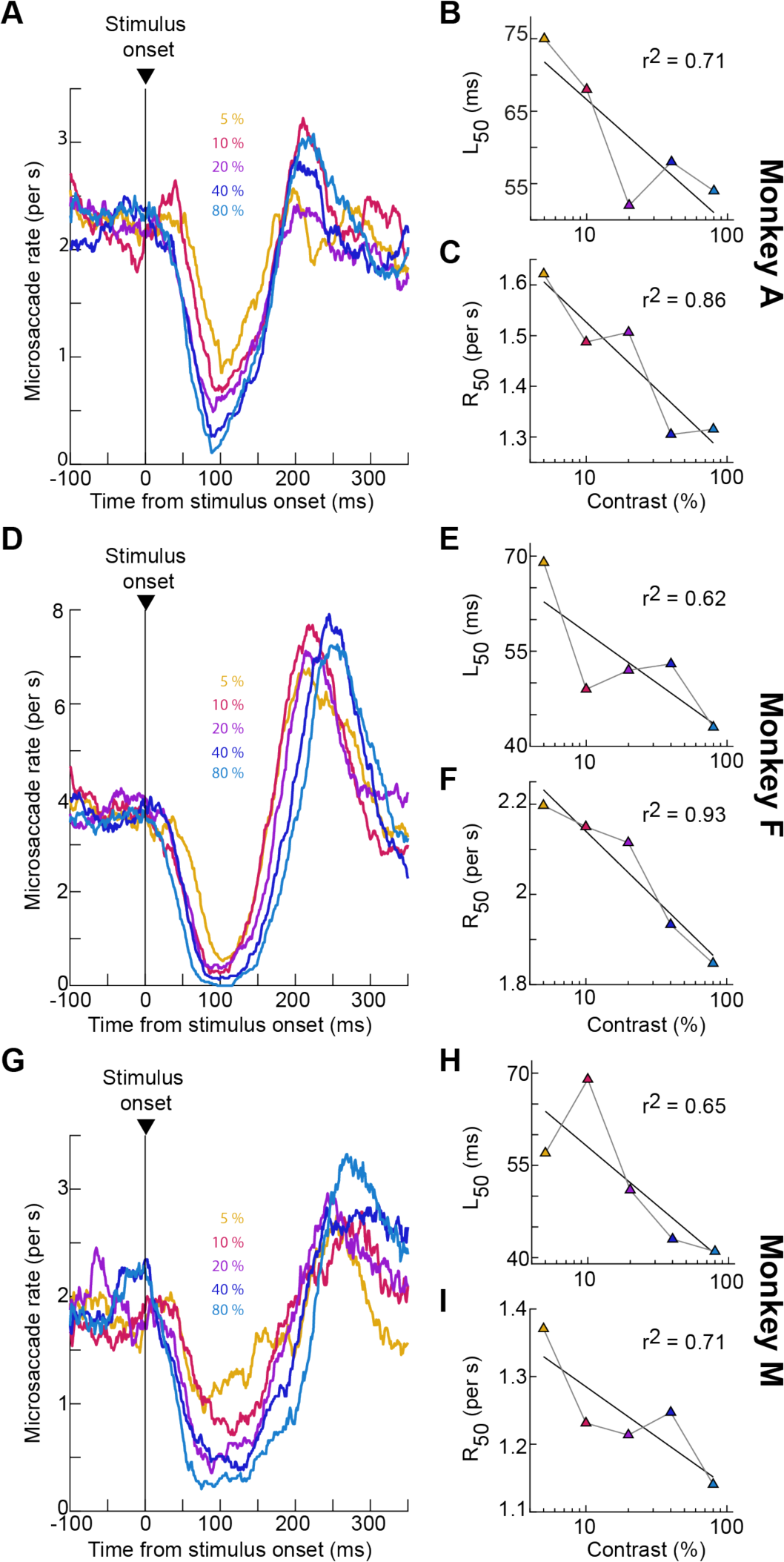
Earlier saccadic inhibition with higher contrasts of large stimuli. **(A-C)** Similar analyses to those in Figs. 2, 3, but now relating saccadic inhibition in monkey A to the contrast of a full-screen flash appearing. Both saccadic inhibition time (L_50_) and strength (R_50_) were contrast-dependent: L_50_ decreased with increasing contrast, and inhibition strength increased with increasing contrast (evidenced by reduced R_50_ rates). **(D-F)** Similar observations for monkey F. **(G-I)** Similar observations for monkey M.

We next ran another experiment in which stimulus contrast was again manipulated. However, in this case, the stimulus onset consisted of a small disc of radius 0.51 deg (29). This disc appeared on the display and remained on for a few hundred milliseconds, but it was to be ignored by the monkeys. The location of the disc varied from session to session, especially because the data from this experiment came from a previous neurophysiological study in which we were also recording superior colliculus visual neural activity (29). Here, we analyzed the negative luminance polarity conditions from that study (to behaviorally match them with the experiment of Fig. 4 using dark contrasts). As can be seen from Fig. 5, even with small, localized stimuli, L_50_ still decreased with increasing stimulus contrast, consistent with the results of Fig. 4.

**Figure 5.**
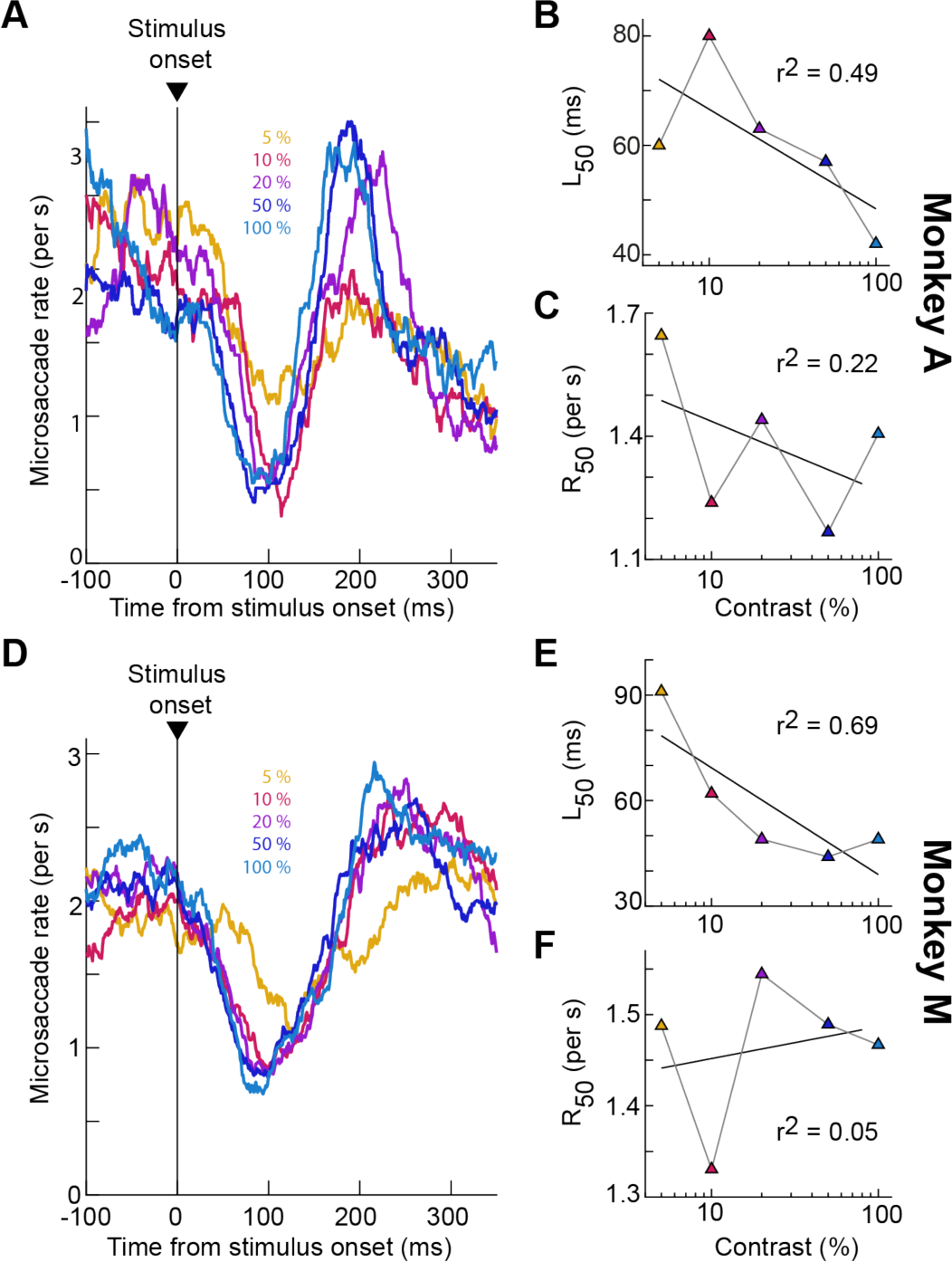
Earlier saccadic inhibition with higher contrasts of small, localized stimuli away from the oculomotor goals of ongoing saccades. **(A-C)** Similar analyses to Fig. 4, but now with the stimulus being a small disc (radius 0.51 deg) appearing somewhere on the display away from where the ongoing saccades were being generated. Saccadic inhibition time (L_50_) still decreased with increasing contrast. The rate effect (R_50_) was less clear as in Fig. 4, likely because the stimulus onset was actively ignored (also see Fig. 6 for additional evidence). **(D-F)** Similar observations from monkey M. Here, the rate effect was even weaker than in monkey A (also see Fig. 6).

The R_50_ effects were noisier in Fig. 5, only showing a more convincing negative trend for monkey A. This could be because the saccadic inhibition effect was overall weaker in this experiment than in the experiment of Fig. 4, which is itself consistent with the size tuning results of Figs. 1-2 above. That is, in Fig. 5, the minimum saccade rate that was reached during peak saccadic inhibition was higher than that in Fig. 4, an observation that is at least partially due to the smaller stimulus sizes (Figs. 1-2). For example, at 100% contrast, the minimum saccade rate in monkeys A, F, and M was 0.15, 0, and 0.33 saccades/s, respectively, in Fig. 4; it was 1.4 and 1.47 saccades/s in monkeys A and M, respectively, in Fig. 5. This led us to next ask what happens if the stimulus onset was to be foveated as opposed to be completely ignored.

#### Stronger saccadic inhibition when appearing stimuli are targets for foveation

The results of Figs. 4, 5 demonstrate that saccadic inhibition depends on stimulus contrast in general, but that an ignored small stimulus away from the oculomotor targets of the ongoing saccadic activity may be associated with generally weaker peak inhibition than a larger visual transient spanning the retinotopic target locations of the ensuing saccades (since the transient covered the fixation spot). That is, at the time of peak saccadic inhibition, there was still a higher likelihood of saccade occurrence with the ignored eccentric stimulus (Fig. 5) than with a large visual transient covering the peri-fovea (where our small saccades were targeted in our gaze fixation tasks) (Fig. 4). However, if the eccentric stimulus is now to be foveated, then the interruption by the visual onset (1) should eventually lead to a foveating eye movement towards the stimulus. In this case, post-inhibition saccades are much more cognitively controlled; they are targeted eye movements towards the appearing stimuli. We found that in this case, saccadic inhibition became all-or-none. Specifically, we repeated the same experiment of Fig. 5, but now requiring the monkeys to foveate the appearing stimulus.

Saccadic inhibition as caused by a visual onset in this new foveating eye movement experiment generally followed a similar timeline to saccadic inhibition when the appearing stimulus was completely ignored in the previous experiment. For example, Fig. 6A, C, E shows the saccade rate curves around stimulus onset from the 100% contrast condition in the two cases. The black rate curves replicate the 100% contrast data from Fig. 5, and they are included in Fig. 6 only up to the peak inhibition time. The blue rate curves instead show saccade rate when the task was to foveate the appearing target after the stimulus-driven saccadic inhibition had begun. In this case, we plotted the rate curves until the time of the foveating saccade that had the lowest reaction time from stimulus onset. Note also that monkey F only performed the foveating saccade version of the task, so we did not show any black curve in this monkey’s panel. As can be seen, in all three animals, when the goal was to foveate the appearing eccentric stimulus, saccadic inhibition was an all-or-none phenomenon (that is, saccade rate dropped to zero). While it is true that the baseline (pre- stimulus) saccade rate was different in the two tasks, peak saccadic inhibition in Fig. 5 never caused zero saccade rates during the inhibition period, even at high contrast (the black curves in Fig. 6 are truncated at the minimum saccade rate and were always well above zero). Consistent with this, across all contrasts, the R_50_ measure in all three animals was lower than the same measure from the very similar task of Fig. 5. This comparison between the two tasks is rendered easier in Fig. 6B, D, F, plotting R_50_ from the data of Fig. 5 in the same panels as R_50_ from this additional experiment (again, monkey F was not tested in the fixation version of the task, so only the current experiment’s results are shown; also, monkey M was not tested with all contrasts in this experiment, so only the tested data points are shown). R_50_ was lower in the current experiment than in the previous one.

**Figure 6.**
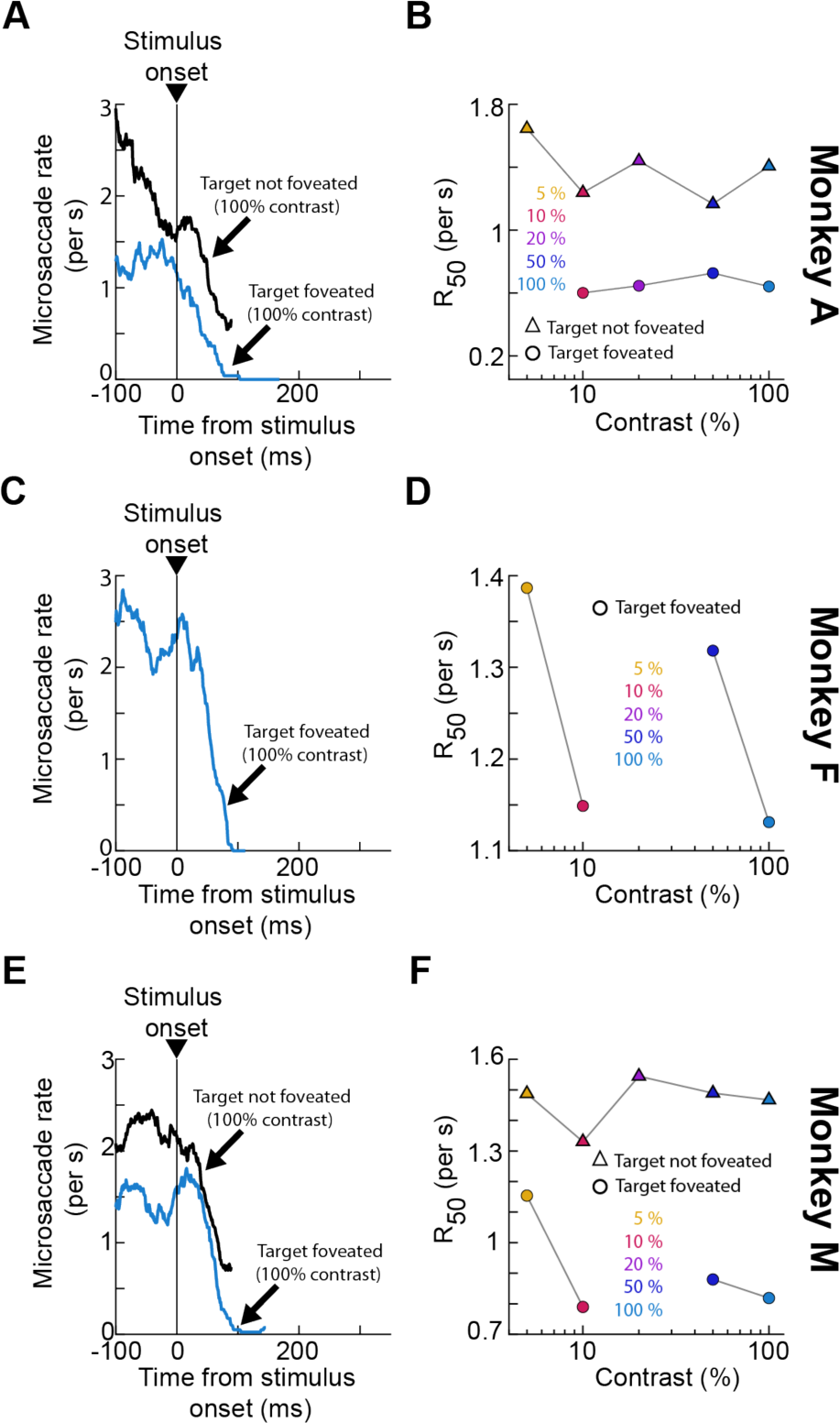
Stronger saccadic inhibition when the appearing stimuli were to be later foveated. **(A)** The black curve shows the same saccade rate curve as that from Fig. 5 for 100% contrast stimuli in monkey A. The blue curve shows the saccade rate curve for the same stimulus and monkey, but now when the stimulus was to be subsequently foveated with a targeting eye movement (Experiment 5). The black curve is truncated at the point of maximum saccadic inhibition, and the blue curve is truncated at the time of the shortest latency foveating saccade. Saccadic inhibition started at approximately the same time in both cases (the black curve had a time varying pre-stimulus saccade rate in this monkey as we also saw in Fig. 3A; this monkey tended to perform fixation tasks by gradually decreasing saccade rate in anticipation of stimulus onset and trial end). However, saccadic inhibition was all-or-none when a subsequent foveating saccade was made. **(B)** Consistent with this, across all tested contrasts in both experiments, R_50_ was lower in the foveated target condition. Note that the data for the condition without foveating saccades shown here is the same as that in Fig. 5C (included here for easier comparison to the other curve). **(C, D)** Similar analyses for monkey F. This monkey did not perform the experiment of Fig. 5, but the data from the current experiment still show all-or-none saccadic inhibition, consistent with monkey A. **(E, F)** Similar analyses for monkey M. Here, both variants of the task were collected, and the same results as with monkey A can be seen. That is, inhibition time was similar in both task variants; however, when the appearing target was later foveated, saccadic inhibition magnitude was much stronger (smaller R_50_).

Therefore, saccadic inhibition can have generally similar time courses depending on the subsequent post-inhibition oculomotor behavior (Fig. 6A, C, E), but the peak inhibition strongly depends on such behavior. Naturally, in this version of the task, the fixation spot was also extinguished at the same time as when the eccentric stimulus appeared. Since the active oculomotor behavior was generally aimed at the fixation spot (18, 39, 40), it could be that we obtained stronger saccadic inhibition in this case because there was a double visual transient (a peripheral target onset as well as a foveal target offset). Nonetheless, these data touch on an interesting question about how multiple different orienting behaviors can be coordinated around the time of stimulus onsets, and they can inform neurophysiological studies of both saccade generation and fixation maintenance in the face of asynchronous external inputs (1).

### Dependence of saccadic inhibition on motion direction

In our final experiment, the stimulus onset consisted of a drifting grating possessing one of eight possible motion directions and one of two possible temporal frequencies. We first analyzed the influence of motion direction on saccadic inhibition, by pooling across temporal frequencies. Figure 7A shows example saccade rate curves around the time of the onset of the drifting gratings. The top panel shows saccade rate from monkey A when the grating was drifting upwards, and the middle panel shows saccade rate when the grating was drifting leftwards. The bottom panel describes saccade rate with downward drifting gratings. In all cases, the location of the stimulus was the same; only the motion direction of the gratings was different across panels. Saccadic inhibition occurred earlier for the horizontal motion direction than for the vertical motion directions (vertical, colored lines indicate L_50_ for each case). Figure 7D, G shows similar observations across all three monkeys.

**Figure 7.**
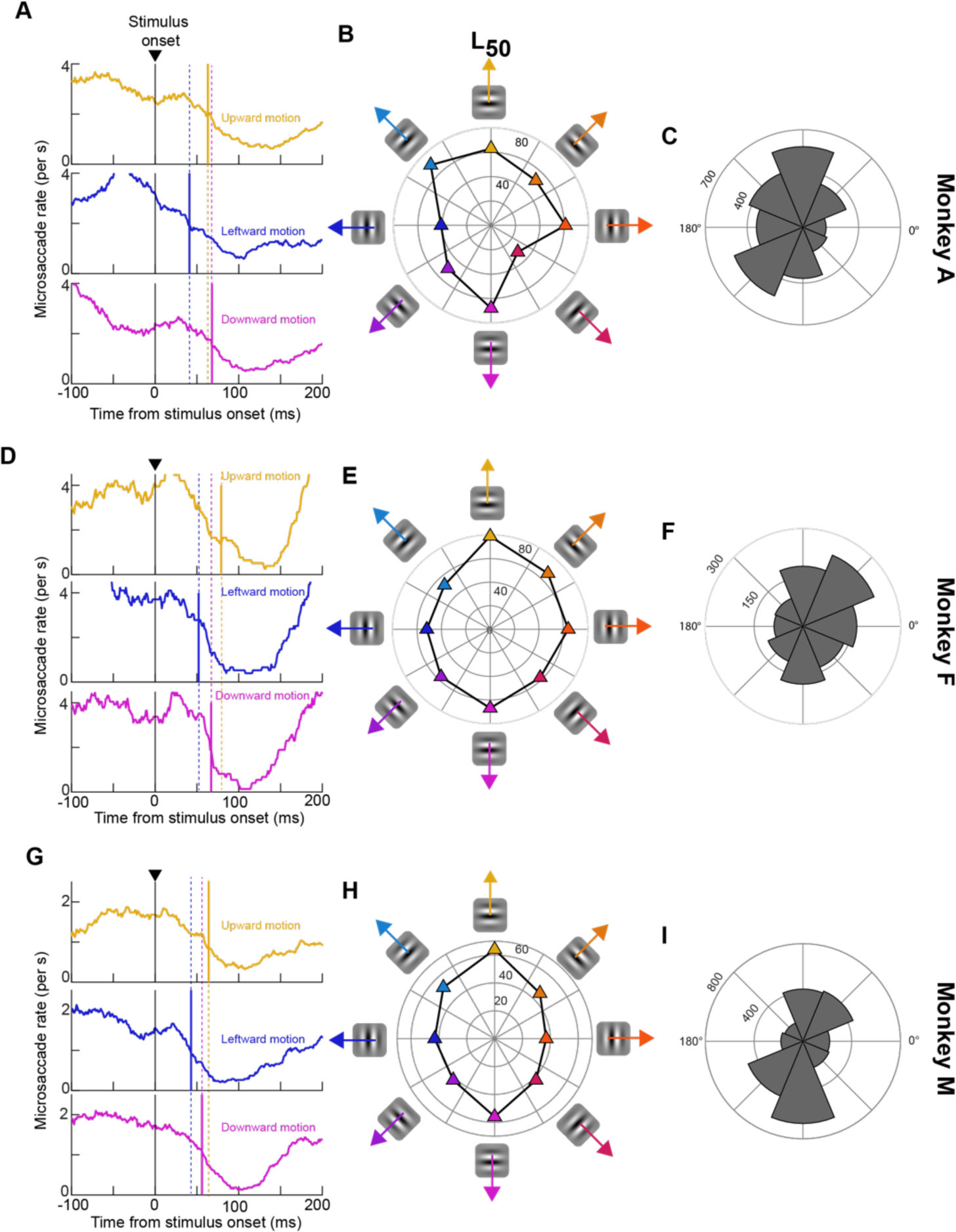
Later saccadic inhibition for vertical motion directions. **(A)** Saccade rate curves of monkey A from three example motion directions in Experiment 6. Each curve is truncated vertically and horizontally to focus on the saccadic inhibition phase. The vertical colored lines indicate L_50_ for their respective saccade rate curves. As can be seen, saccadic inhibition occurred earlier for leftward motion directions than for both upward and downward motion directions. **(B)** Values of L_50_ in this experiment and animal for all tested motion directions. Horizontal motions generally had shorter L_50_ values than vertical directions. Up-left motion directions also had the longest L_50_ values. **(C)** We plotted the angular distribution of saccade directions during a pre-stimulus baseline interval and noticed that the biases in **B** could be correlated with those in the current panel. For example, the monkey made more saccades in the upward and leftward direction, and up-left motions were associated with delayed saccadic inhibition. **(D-F)** Similar observations in monkey F. Again, horizontal motion directions were associated with smaller L_50_ values than vertical motion directions. Moreover, in this case, L_50_ was additionally longer in the up-right than in the up-left motion direction (**E**), and this was correlated with the monkey’s intrinsic bias to make more baseline saccades towards the upper right direction (**F**). **(G-I)** Similar observations in monkey M. Again, horizontal motion directions were consistently associated with earlier saccadic inhibition times than vertical motion directions.

Interestingly, when we tested all motion directions in each animal (Fig. 7B, E, H), we found slightly variable dependencies of the time of saccadic inhibition on motion direction in each individual. Specifically, while it was generally true that horizontal motion directions were associated with earlier L_50_ times than vertical ones (Fig. 7A, D, G), each monkey showed a specific set of additional motion directions with particularly long L_50_ times relative to the others. In monkey A, this was the case for upward-leftward motion directions; in monkey F, this was the case for upward-rightward motion directions; and in monkey M, this was the case for upward or downward motion directions.

We next considered a potential correlate of such individual monkey idiosyncrasy. Specifically, we analyzed the direction distribution of microsaccades in each monkey during baseline fixation, before any stimulus appeared. To do this, we picked an interval before stimulus onset (final 100 ms before the onset event occurred) across all motion directions. We then plotted the angular distribution of saccade directions in such baseline interval. The saccade direction distributions of all three animals are shown in Fig. 7C, F, I. As can be seen, the dependence of L_50_ in each animal was correlated with the animal’s intrinsic saccade direction distribution during baseline intervals. For example, monkey A tended to make more up-left oblique saccades, whereas monkey F tended to make more up-right oblique movements. In both cases, L_50_ was longer in a corresponding direction for the respective animal. Similarly, monkey M made more frequent vertical saccades than horizontal saccades (Fig. 7I), and this again was correlated with longer L_50_ times for vertical motion directions.

While this is just a correlation, these observations might suggest that each monkey experiences more frequent retinotopic motion directions during self-movement because of the intrinsic biases in saccade directions. It could, therefore, be that saccadic inhibition is easier for the more-frequently experienced retinal motion directions. Consistent with this, L_50_ was shorter for downward-rightward than upward-leftward motion directions in monkey A, and this monkey tended to make more up-left saccades in general (experiencing more downward-rightward retinal image shifts during eye movements). In monkey F, L_50_ was shorter for leftward and leftward-upward motion directions, which are the opposite of the more frequent saccade directions in this animal. This hypothesis is not quite clear for monkey M, but this monkey was equally likely to make upward or downward vertical saccades, potentially balancing these motion directions in his retinal-image shift experience across eye movements.

In any case, it is interesting to observe a dependence of saccadic inhibition on motion direction in our experiments. It is also interesting that horizontal motions were generally easier to inhibit (shorter L_50_ times) than vertical motions. This could reflect stronger visual signals for horizontal motion directions, and it could fit with a relatively large literature showing how vision in the horizontal cardinal dimension might be better than vision in the vertical cardinal dimension (52–54).

### Dependence of saccadic inhibition on motion speed

Finally, we pooled across all motion directions from Fig. 7 to check whether there was an impact of motion speed on saccadic inhibition. We had two motion speeds in the drifting gratings, characterized by two different temporal frequencies. We found that saccadic inhibition timing in all three monkeys generally did not strongly depend on motion speed (Fig. 8; colored L_50_ lines). However, faster speeds caused a deeper and longer lasting minimum of saccade rate than slower speeds in all three animals (Fig. 8). Thus, recovery from saccadic inhibition was harder for the faster motion speeds in all three monkeys.

**Figure 8.**
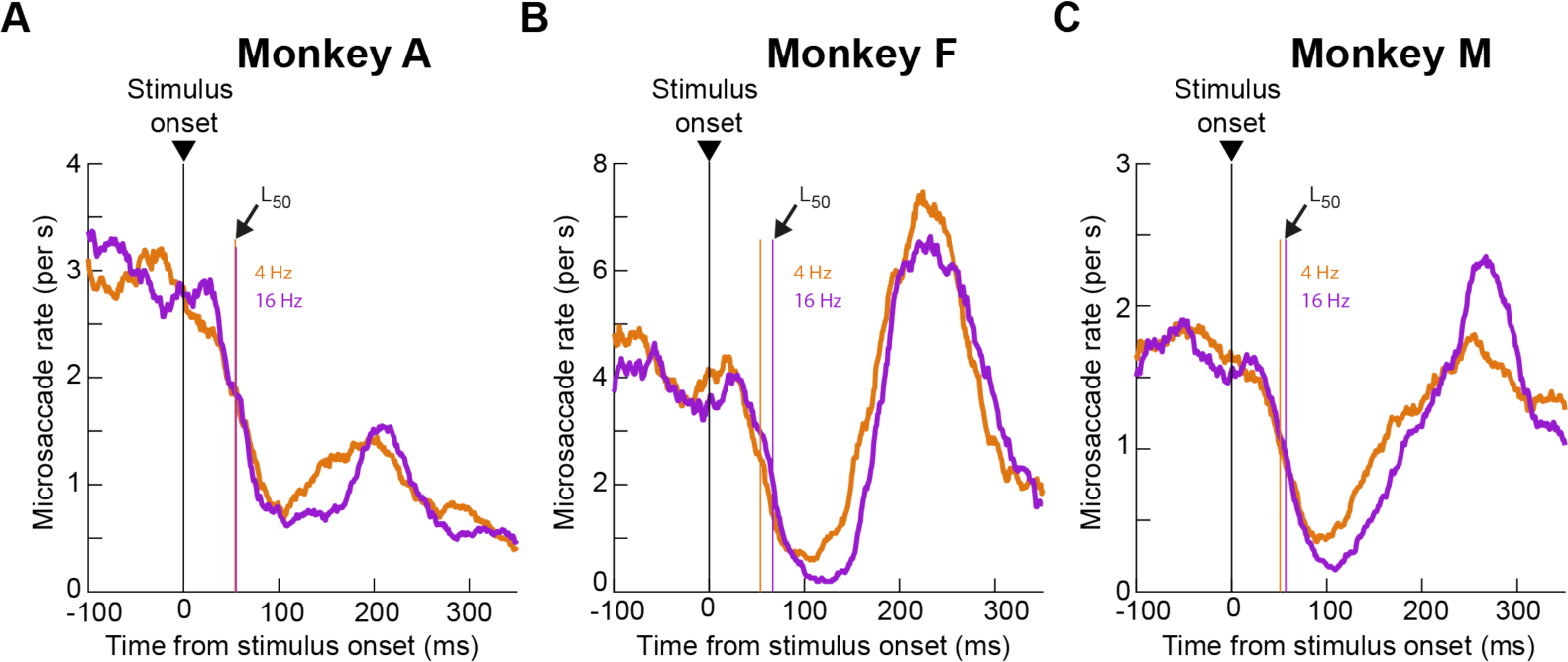
Longer lasting saccadic inhibition for faster motions. **(A-C)** For each monkey, we collapsed across all motion directions from Experiment 6, and we plotted saccade rate curves. Saccadic inhibition timing (vertical colored lines) was generally similar for different motion speeds (caused by the different temporal frequencies of the drifting gratings). However, in all cases, the faster speed was associated with a longer lasting saccadic inhibition period before the subsequent post-inhibition saccades.

## Discussion

We characterized the properties of saccadic inhibition in rhesus macaque monkeys as a function of different visual feature dimensions. We found that saccadic inhibition in these animals systematically depends on stimulus size, spatial frequency, contrast, and motion direction. We also found that if appearing stimuli are subsequently foveated as opposed to being ignored, saccadic inhibition is stronger and becomes much more like an all-or-none phenomenon. On the other hand, relatively small eccentric “distractors” that are ignored have significantly milder inhibitory effects on saccade generation.

Some of the feature dimensions that we tested, like stimulus contrast, were also tested previously in humans (9–11). The similarity of our findings in monkeys to those observations in humans reinforces our belief that macaque monkeys are a suitable model system for exploring the neural mechanisms of saccadic inhibition. In fact, recent transcranial magnetic stimulus (TMS) studies of the human homolog of the monkey frontal eye fields also affirm the utility of monkeys for investigating neural mechanisms of phenomena related to saccadic inhibition (47, 48). Specifically, these TMS studies disrupted post-inhibition saccades with disruption of frontal eye field activity, consistent with the predictions from reversible inactivation of the frontal eye fields of macaque monkeys (26). This homology between the two species is exactly why we performed the current experiments. These experiments provide, in our view, a reference frame with which we hope to inform our upcoming neurophysiological studies of saccadic inhibition in the near future.

We think that it is likely to see future neurophysiological experiments revealing an important role for oculomotor control circuits in the midbrain and brainstem in mediating saccadic inhibition. Indeed, visual responses in the superior colliculus already hint that such responses in oculomotor control circuitry can matter a great deal for saccade generation. For example, collicular visual responses occur earlier for low rather than high spatial frequency stimuli, and this mimics the patterns of saccadic reaction times in visually-guided saccade paradigms (50). Similarly, express saccades (saccades with reaction times less than around 90-100 ms) seem to be triggered by direct readout of the spatial locus of superior colliculus visual bursts (occurring within 50-100 ms from stimulus onset) (55). Thus, in the case of express saccades, visual sensory responses do indeed have a privileged and direct impact on saccade generation. Likewise, we think that visual responses in the oculomotor control network should have a privileged and direct impact on saccadic inhibition, again because of the very short latency with which inhibition is achieved. In this case, we might predict (1, 20) that such an impact of visual responses should be inhibitory (rather than excitatory as in the case of the superior colliculus and express saccades). Such an inhibitory effect can arise if omnipause neurons in the brainstem (56, 57) exhibited visual pattern responses to stimulus onsets of different feature properties, and if these responses were consistent with the feature tunings that we discovered in the current study.

The above thoughts lead to the idea that the scene analysis that takes place by oculomotor control circuits in the brain, via the sensitivity of these circuits to visual inputs, is a reformatted representation of the scene. That is, it may not be needed for the superior colliculus and other oculomotor control circuits to just inherit the visual properties of the primary visual cortex or other cortical areas, even if the signals eventually come from these cortical areas. Rather, the representation is reformatted for something useful for the oculomotor system (28). This is not unlike evidence that the superior colliculus seems to favor the upper visual field (58) when ventral stream visual cortical areas might favor the lower visual field (59). Thus, the oculomotor system “sees” a filtered representation of the visual scene that is not necessarily the same as what cortical areas for scene analysis and interpretation might “see”, and this is still the case even if it is the signals in the early cortical visual areas (like primary visual cortex) that are ultimately relayed to the oculomotor control network.

If that is indeed the case, then one question that might arise in relation to our results in the current article could be: why would the oculomotor system need stronger and earlier saccadic inhibition for low spatial frequencies? One possibility is that low spatial frequency stimuli are quite salient, and excitatory structures like the superior colliculus already favor these stimuli (50). Thus, because any spike in the superior colliculus can have an excitatory impact on the oculomotor system (55, 60), the inhibitory system that balances coordination with exogenous stimuli (1) would need to be equally potent for low spatial frequency stimuli. A similar kind of logic also applies for stimulus contrast and size. Thus, we anticipate that circuits driving saccadic inhibition should have similar feature tuning preferences to circuits, such as the superior colliculus, that drive saccade generation.

We also find the motion direction effects on saccadic inhibition particularly intriguing. In all three animals, we found earlier saccadic inhibition, as evidenced by smaller L_50_ values, for horizontal rather than vertical motion directions. This is interesting from the perspective of visual field asymmetries and oculomotor behavior, including in short-term memory (52, 61). In Results, we also framed this anisotropy as potentially being related to the baseline anisotropies of saccade generation in the individual animals. However, both explanations may not necessarily be mutually exclusive. For example, it could be that individual saccade directions are more or less likely in one animal exactly because of the animal’s specific instantiation of visual field anisotropies in neural circuits. Indeed, given that saccades during fixation of a target (as in our experiments) primarily correct eye position errors even in explicit cueing tasks (40), the biased distributions of saccades in individual animals might reflect biased distributions of drift eye movements in the animals. Such drift eye movements, and the related saccades that intersperse them, continually expose the visual system to specific patterns of retinal image motion. They could thus either reflect or shape individual visual representational anisotropies in a given animal. It would be interesting in the future to relate saccadic inhibition properties to explicit experimentally controlled retinal image drifts.

In our stimulus contrast experiments, we also investigated the case in which a small stimulus was more like a distractor, or whether it became behaviorally relevant by requiring its foveation after saccadic inhibition was completed. We found that saccade rate dropped down to zero in the latter case. This makes functional sense. Every saccade is a bottleneck, and no other saccade can be generated at the same time. Therefore, for the target to be foveated, saccade rate had to drop to zero. However, post-inhibition saccades in the distractor case are also a bottleneck, and it is interesting to contemplate why saccadic inhibition was weaker in this case. One possibility is that there were two sensory transients in the foveating condition: in addition to the stimulus onset, the fixation spot was removed simultaneously to instruct the animals about the behavioral relevance of the appearing stimulus (and that it should be foveated). Therefore, it could be that there was a larger sensory drive for the inhibitory circuits. It would be interesting in the future to study additional top-down impacts on saccadic inhibition, but from a neurophysiological perspective to exploit the use of monkeys as a model system for the phenomenon.

Finally, several of our experiments included large stimulus onsets (e.g full-screen flashes). We recently found that such onsets are associated with a stimulus-driven tiny drift of eye position (much smaller than microsaccades) (62). Such a drift response seems to also be stimulus driven. However, the detailed feature tuning properties of this response, like in the case of saccadic inhibition, are still not fully explored. Given that the drift response seems to be coordinated with the time of saccadic inhibition, our goal in the near future is to document the feature tuning properties of the drift response in more detail, like we did here for saccadic inhibition. This way, we would have a rich behavioral characterization of oculomotor phenomena related to the coordination between internal active perceptual state and asynchronous exogenous stimuli. Such characterization should open the door for interesting new insights about the underlying brain mechanisms of active perception and cognition.

## Data availability

Table 1 in the Appendix lists all relevant summary statistics. Raw data will be made available upon request.

## Grants

We were funded by the following grants from the Deutsche Forschungsgemeinschaft (DFG; German Research Foundation): BU4031/1-1, HA6749/4-1, BO5681/1-1, HA6749/3-1, and SFB 1233, Robust Vision: Inference Principles and Neural Mechanisms, TP11, project number: 276693517.

## Appendix

**Table 1.** Numerical measurements included in this study. Baseline saccade rate was obtained from the final 100 ms of fixation before stimulus onset. The Materials and Methods section describes how L_50_ and R_50_ were obtained.

| Experiment | Animal | Baseline saccade rate (and 95% confidence interval) (saccades/s) | Condition | $L_{50}$ (and 95% confidence interval) (ms) | $R_{50}$ (and 95% confidence interval) (saccades/s) | Number of trials |
| --- | --- | --- | --- | --- | --- | --- |
| 1: Size tuning | A | 2.00 (1.90, 2.17) | 0.09 deg | 54 (47, 63) | 1.43 (1.26, 1.63) | 1371 |
|  |  |  | 0.18 deg | 60 (45, 66) | 1.27 (1.09, 1.49) | 1344 |
|  |  |  | 0.36 deg | 56 (51, 62) | 1.08 (0.92, 1.21) | 1388 |
|  |  |  | 0.72 deg | 55 (43, 60) | 1.10 (0.10, 1.25) | 1402 |
|  |  |  | 1.14 deg | 47 (40, 53) | 1.25 (1.15, 1.45) | 1341 |
|  |  |  | 2.28 deg | 48 (38, 57) | 1.14 (1.07, 1.23) | 1447 |
|  |  |  | 4.56 deg | 49 (43, 57) | 1.04 (0.87, 1.17) | 1369 |
|  |  |  | 9.12 deg | 50 (44, 55) | 1.02 (0.89, 1.07) | 1394 |
|  | F | 3.49 (3.35, 3.64) | 0.09 deg | 75 (67, 80) | 2.17 (2.06, 2.41) | 1033 |
|  |  |  | 0.18 deg | 65 (62, 71) | 1.98 (1.76, 2.11) | 999 |
|  |  |  | 0.36 deg | 55 (50, 60) | 1.96 (1.78, 2.04) | 1017 |
|  |  |  | 0.72 deg | 61 (59, 65) | 1.77 (1.68, 1.91) | 1044 |
|  |  |  | 1.14 deg | 55 (51, 58) | 1.76 (1.56, 1.95) | 1093 |
|  |  |  | 2.28 deg | 49 (42, 50) | 1.81 (1.69, 1.94) | 1089 |
|  |  |  | 4.56 deg | 45 (43, 47) | 1.90 (1.76, 1.97) | 1003 |
|  |  |  | 9.12 deg | 43 (38, 44) | 1.83 (1.74, 1.97) | 1068 |
|  | M | 1.90 (1.74, 2.06) | 0.09 deg | 76 (66, 99) | 1.21 (0.99, 1.24) | 628 |
|  |  |  | 0.18 deg | 59 (32, 68) | 1.08 (0.92, 1.60) | 655 |
|  |  |  | 0.36 deg | 54 (45, 61) | 0.94 (0.78, 1.09) | 661 |
|  |  |  | 0.72 deg | 42 (29, 47) | 1.01 (0.91, 1.12) | 710 |
|  |  |  | 1.14 deg | 40 (31, 46) | 0.86 (0.72, 0.98) | 713 |
|  |  |  | 2.28 deg | 50 (41, 57) | 0.91 (0.81, 1.06) | 660 |
|  |  |  | 4.56 deg | 44 (36, 51) | 1.01 (0.89, 1.08) | 664 |
|  |  |  | 9.12 deg | 39 (25, 48) | 0.93 (0.80, 1.40) | 632 |
| 2: Spatial frequency | A | 2.62 (2.42, 2.83) | 0.5 cpd | 55 (41, 67) | 1.93 (1.51, 2.15) | 483 |
|  |  |  | 1 cpd | 56 (42, 69) | 1.66 (1.33, 1.89) | 484 |
|  |  |  | 2 cpd | 64 (49, 74) | 1.80 (1.45, 2.08) | 484 |
|  |  |  | 4 cpd | 68 (39, 91) | 2.05 (1.59, 2.26) | 487 |
|  |  |  | 8 cpd | 83 (63, 95) | 1.75 (1.46, 2.05) | 484 |
|  | F | 3.60 (3.36, 3.85) | 0.5 cpd | 61 (48, 66) | 1.84 (1.61, 2.26) | 385 |
|  |  |  | 1 cpd | 62 (53, 70) | 2.14 (1.87, 2.42) | 389 |
|  |  |  | 2 cpd | 58 (39, 71) | 2.45 (2.05, 2.82) | 380 |
|  |  |  | 4 cpd | 81 (56, 89) | 2.32 (2.06, 2.75) | 385 |
|  |  |  | 8 cpd | 75 (62, 88) | 2.46 (1.98, 2.71) | 383 |
| **3: Contrast (full screen)** | A | 2.34 (2.22, 2.46) | 5% | 75 (65, 86) | 1.62 (1.39, 1.76) | 1315 |
|  |  |  | 10% | 68 (61, 77) | 1.49 (1.27, 1.63) | 1308 |
|  |  |  | 20% | 52 (43, 62) | 1.51 (1.26, 1.61) | 1315 |
|  |  |  | 40% | 58 (50, 66) | 1.30 (1.13, 1.44) | 1321 |
|  |  |  | 80% | 54 (45, 60) | 1.31 (1.12, 1.45) | 1321 |
|  | F | 3.58 (3.40, 3.75) | 5% | 69 (58, 75) | 2.20 (1.92, 2.44) | 767 |
|  |  |  | 10% | 49 (37, 58) | 2.15 (1.84, 2.38) | 771 |
|  |  |  | 20% | 52 (45, 59) | 2.11 (1.90, 2.30) | 766 |
|  |  |  | 40% | 53 (47, 59) | 1.93 (1.69, 2.16) | 776 |
|  |  |  | 80% | 43 (36, 50) | 1.84 (1.62, 2.03) | 760 |
|  | M | 1.94 (1.8, 2.09) | 5% | 57 (46, 104) | 1.37 (1.15, 1.55) | 785 |
|  |  |  | 10% | 69 (55, 99) | 1.23 (0.96, 1.53) | 789 |
|  |  |  | 20% | 51 (34, 61) | 1.21 (1.00, 1.39) | 790 |
|  |  |  | 40% | 43 (30, 63) | 1.25 (0.93, 1.38) | 785 |
|  |  |  | 80% | 41 (32, 49) | 1.14 (0.95, 1.28) | 788 |
| **4: Contrast (small stimulus)** | A | 2.15 (1.98, 2.32) | 5% | 60 (44, 102) | 1.64 (1.32, 1.87) | 627 |
|  |  |  | 10% | 80 (54, 90) | 1.24 (1.03, 1.52) | 623 |
|  |  |  | 20% | 63 (47, 77) | 1.44 (1.16, 1.61) | 622 |
|  |  |  | 50% | 57 (43, 68) | 1.17 (0.95, 1.35) | 626 |
|  |  |  | 100% | 42 (28, 58) | 1.40 (1.06, 1.57) | 623 |
|  | M | 2.21 (2.10, 2.33) | 5% | 91 (71, 111) | 1.49 (1.32, 1.66) | 1689 |
|  |  |  | 10% | 62 (41, 78) | 1.33 (1.20, 1.50) | 1689 |
|  |  |  | 20% | 49 (32, 65) | 1.54 (1.34, 1.65) | 1682 |
|  |  |  | 50% | 44 (29, 58) | 1.49 (1.32, 1.60) | 1677 |
|  |  |  | 100% | 49 (39, 59) | 1.47 (1.32, 1.61) | 1692 |
| **5: Contrast (foveating saccade)** | A | 1.29 (1.14, 1.44) | 10% | 60 (43, 70) | 0.60 (0.48, 0.88) | 498 |
|  |  |  | 20% | 60 (40, 70) | 0.64 (0.50, 0.87) | 439 |
|  |  |  | 50% | 53 (35, 67) | 0.72 (0.57, 0.91) | 495 |
|  |  |  | 100% | 38 (20, 54) | 0.64 (0.47, 0.83) | 497 |
|  | F | 2.40 (2.25, 2.55) | 5% | 71 (62, 78) | 1.39 (1.19, 1.55) | 915 |
|  |  |  | 10% | 72 (63, 78) | 1.15 (1.00, 1.32) | 878 |
|  |  |  | 50% | 59 (52, 66) | 1.32 (1.17, 1.48) | 888 |
|  |  |  | 100% | 58 (49, 66) | 1.13 (0.97, 1.37) | 883 |
|  | M | 1.51 (1.38, 1.64) | 5% | 66 (44, 69) | 1.15 (0.43, 1.92) | 52 |
|  |  |  | 10% | 61 (51, 68) | 0.79 (0.64, 0.93) | 805 |
|  |  |  | 50% | 56 (49, 65) | 0.88 (0.70, 1.01) | 808 |
|  |  |  | 100% | 62 (54, 69) | 0.82 (0.61, 0.93) | 806 |
| 6: Motion direction (pooled across temporal frequencies) | A | 2.78 (2.57, 2.98) | Up | 63 (48, 80) | 2.00 (1.58, 2.26) | 479 |
|  |  |  | Up-right | 52 (37, 73) | 1.57 (1.18, 1.80) | 479 |
|  |  |  | Right | 61 (52, 73) | 1.81 (1.45, 2.03) | 486 |
|  |  |  | Down-right | 31 (19, 53) | 2.24 (1.74, 2.37) | 482 |
|  |  |  | Down | 68 (49, 79) | 1.64 (1.33, 1.92) | 480 |
|  |  |  | Down-left | 50 (37, 63) | 1.66 (1.36, 1.91) | 484 |
|  |  |  | Left | 41 (21, 59) | 2.09 (1.71, 2.36) | 479 |
|  |  |  | Up-left | 70 (56, 84) | 1.80 (1.42, 2.04) | 486 |
|  | F | 4.37 (3.96, 4.77) | Up | 79 (52, 100) | 1.64 (1.37, 2.48) | 157 |
|  |  |  | Up-right | 67 (44, 77) | 2.23 (1.70, 2.82) | 154 |
|  |  |  | Right | 64 (48, 76) | 2.48 (1.75, 2.82) | 150 |
|  |  |  | Down-right | 58 (44, 70) | 2.11 (1.58, 2.56) | 154 |
|  |  |  | Down | 67 (51, 80) | 2.01 (1.45, 2.60) | 148 |
|  |  |  | Down-left | 57 (32, 77) | 2.40 (1.88, 2.88) | 153 |
|  |  |  | Left | 52 (33, 69) | 2.70 (1.90, 2.98) | 151 |
|  |  |  | Up-left | 53 (29, 67) | 2.24 (1.78, 2.90) | 151 |
|  | M | 2.0 (1.84, 2.15) | Up | 64 (44, 73) | 0.92 (0.78, 1.16) | 737 |
|  |  |  | Up-right | 46 (25, 58) | 0.98 (0.81, 1.19) | 732 |
|  |  |  | Right | 37 (26, 50) | 0.99 (0.73, 1.12) | 745 |
|  |  |  | Down-right | 42 (25, 56) | 0.97 (0.78, 1.11) | 745 |
|  |  |  | Down | 56 (40, 68) | 1.07 (0.84, 1.25) | 752 |
|  |  |  | Down-left | 42 (30, 54) | 0.95 (0.75, 1.14) | 753 |
|  |  |  | Left | 43 (33, 54) | 1.01 (0.81, 1.20) | 756 |
|  |  |  | Up-left | 52 (39, 59) | 1.03 (0.86, 1.24) | 741 |

